# Liver biopsies obtained throughout SIV infection reveal evolving interferon stimulated protein expression within distinct monocyte/macrophage subsets

**DOI:** 10.1101/2025.05.01.651668

**Authors:** Nina Derby, Brooke I. Johnson, Sreya Biswas, Miranda Fischer, Katherine A. Fancher, Cristina Luevano-Santos, Sofiya Yusova, Kimberly A. Meyer, Yohannes M. Abraham, Savannah S. Lutz, Cole Fisher, Maria Cristina Pacheco, Jeremy V. Smedley, Benjamin J. Burwitz, Donald L. Sodora

**Affiliations:** Seattle Children’s Research Institute; Oregon National Primate Research Center; Seattle Children’s Hospital; Vaccine and Gene Therapy Institute; University of Louisiana at Lafayette, New Iberia Research Center; University of Washington, Dept of Pediatrics

## Abstract

Liver dysfunction is more common and more severe among people with HIV than in the general population. Our previous transcriptomic analysis identified an increased hepatic type-1 interferon (IFN-1) response in simian immunodeficiency virus (SIV) infected rhesus macaques, including pronounced upregulation of the IFN-1 stimulated gene product MX1. Here, we interrogated the role of different cell types in the IFN-1 response, focusing on macrophage subsets. During the acute phase of SIV infection, the expression of MX1 in liver biopsies was elevated in the resident macrophage Kupffer cells (KCs, CD163+ CD206+) as well as liver sinusoidal endothelial cells (LSECs, CD163– CD206+). Chronic infection was associated with increased MX1 expression both within KCs and the recruited monocyte-derived macrophages (MdMs, CD163+ CD206–) although the KCs were associated with the highest MX1 expression levels. We explored factors in the liver potentially associated with the macrophage IFN-1 response: 1) SIV DNA load, 2) bacterial DNA load (assessed via SIV DNA and 16S bacterial DNA qPCR, respectively), and 3) acute-phase hepatocyte lipid accumulation (steatosis). Both SIV and bacteria were elevated during chronic infection and directly correlated with MdM frequency. KC frequency did not correlate with SIV load but inversely correlated with bacterial load and also correlated directly with hepatocyte microvesicular steatosis in acute infection. Hepatocytes expressed variable levels of MX1 during acute and chronic infection. Overall, our data identify MdMs, LSECs and KCs as contributors to the heightened expression of MX1 in the liver during SIV infection. KCs express the highest levels of MX1 and therefore represent a potential target to reduce liver inflammation in people with HIV.

**AUTHOR SUMMARY:** Liver disease disproportionately affects people with HIV (PWH). However, liver biopsies are generally not indicated in humans, making the immune mechanisms underlying HIV-driven liver dysfunction difficult to study. Using a model of HIV infection in which rhesus macaques are infected with SIV, we previously found a strong upregulation of type 1 interferon (IFN-1) signaling as well as an increased number of CD68+ monocyte/macrophage cells in the liver during chronic infection. Here we used longitudinally collected liver biopsies from SIV-infected macaques to describe the kinetics of the IFN-1 response, focusing on MX1, which was a top differentially expressed gene in our previous transcriptomic screen. We also defined two populations of liver macrophages – resident Kupffer cells and recruited monocyte-derived macrophages – to identify which cells produce MX1, what stimuli the cells respond to, and where the cells accumulate within the liver. Our results indicate that Kupffer cells are the major producers of MX1 during acute and chronic infection and that their frequency in the liver correlates with lipid accumulation (steatosis) during acute infection. These data provide insights into how Kupffer cells may promote inflammation and fibrosis and identify Kupffer cells as an important target to limit liver disease in PWH.

## Introduction

Metabolic dysfunction-associated steatotic liver disease (MASLD) (formerly non-alcoholic fatty liver disease (NAFLD)) has become the most prevalent liver disease worldwide, affecting 25% of the global population including 100 million people in the US (1, 2). In humans, MASLD is diagnosed when more than 5% of hepatocytes contain lipid in the absence of cellular infiltrate (1). The mechanisms that promote MASLD, as well as the more inflammatory metabolic associated steatohepatitis (MASH), are not precisely defined. All four of the major liver cell types – hepatocytes, liver sinusoidal endothelial cells (LSECs), hepatic stellate cells, and macrophages – have been implicated (1, 3, 4). The prevalence of MASLD among people with HIV (PWH) is higher than in the general population, even in the absence of hepatitis virus infection and when HIV replication is effectively suppressed with combination antiretroviral therapy (cART) (5–9). Indeed, a recent U.S. multi-center multi-ethnicity study found steatosis in 49% of the HIV-mono-infected cART-treated subjects sampled (9). Within the setting of HIV infection, not only is MASLD more common than in HIV-uninfected individuals, but it is also more severe with a higher incidence of MASH-defining criteria (10). In PWH, end-stage liver disease is a major cause of non-AIDS mortality (5–8). Currently, no drug therapies to treat MASLD are licensed generally or specifically for PWH, though clinical trials are ongoing (11).

Mounting evidence supports that multiple insults – derived from genetics, metabolism, diet, microbiome and more – induce a hyper-inflammatory and dysfunctional state, including type-1 interferon (IFN-1) signaling, in the livers of people with MASLD (1, 3, 12, 13). While IFN-1 responses are critical in the control of viral infections (14), studies in PWH and in the simian immunodeficiency virus (SIV) rhesus macaque model have shown that the persistence of IFN-1 responses during chronic pathogenic HIV/SIV infections is associated with accelerated disease progression (15, 16). In natural host monkey species that experience a non-pathogenic SIV outcome despite high viral loads, IFN-1 expression resolves during the acute to chronic phase transition, underscoring the benefits of preventing long-term expression of IFN-1s and interferon stimulated genes (ISGs) (17). Numerous studies have documented the roles of inflammation and IFN-1 in the activation of hepatic stellate cells and development of MASH (18). For example, an upregulated IFN-1 response in the livers of obese mice and humans fosters the local expansion of CD8 T cells that promotes metabolic disease (19). In a humanized mouse model of HIV and cART using mice repopulated with human fetal liver, IFN-1 and transforming growth factor-beta (TGF-β) cooperated in the induction of liver fibrosis (20). It is noteworthy that the fibrosis was also associated with the accumulation of M2-skewed macrophages (20).

Foundational studies in the SIV macaque model have documented changes to the hepatic immunological and microbiological landscape during SIV infection (21–31). But as these studies largely focus on the time of necropsy when liver samples can be easily obtained, the results are most often cross-sectional and describe chronic infection. Such work from our laboratory demonstrated that chronic SIV infection induces a heightened IFN-1 response, accumulation of CD68+ monocyte/macrophage cells, and microbial dysbiosis in the liver (30, 31). By employing laparoscopic biopsy collections to evaluate SIV-induced changes to the liver over time, we found that not only are immune cells in the liver impacted by SIV, but hepatocytes are also transiently impacted during acute infection (32). We observed microvesicular steatosis in hepatocytes, increased expression of the peroxisome proliferator activated receptor (*PPARA*, a xenobiotic and lipid sensor) transcript in the liver, and elevated concentrations of aminotransferases and cholesterol in serum (32). Here, using laparoscopic liver biopsies, we assessed the timing of the IFN-1 response in the SIV-infected liver. IFN-1 responsive cells can produce many ISGs, such as MX1, 2’,5’-oligoadenylate synthetase (OAS), ISG15, and various IFITM proteins. We focused on MX1 herein because it is a key ISG upregulated in cells in response to IFN-1 signaling and was among the top upregulated genes in our previous study of the SIV-infected liver (31). We now provide evidence that the IFN-1 response develops during acute infection and is sustained throughout 32 weeks of follow up but evolves over time in the cells involved. Hepatocytes begin to produce MX1 during acute infection and persist throughout chronic infection. Among the liver monocyte/macrophage populations, the IFN-1 response is dominated by macrophages with a Kupffer cell (KC) phenotype that accumulate in the liver in direct correlation with microvesicular steatosis in hepatocytes. In contrast, the monocyte-derived-macrophages (MdMs) produce lower levels of MX1, and correlate with the presence of SIV and bacterial DNA. These results, facilitated by longitudinal sampling, identify distinct roles for different populations of liver macrophages throughout SIV infection and provide key insights into the factors associated with liver inflammation in PWH.

## Results

### Robust IFN-1 response in liver CD68+ cells and hepatocytes in SIV-infected macaques

To evaluate the IFN-1 response within the livers of SIV-infected macaques, expression of the ISG MX1 was assessed by immunofluorescence microscopy in laparoscopically collected liver tissue from 9 SIV-infected and 8 SIV-uninfected (naïve) macaques. Liver biopsies were collected at the baseline 4 weeks prior to SIV infection (Week −4), at Weeks 2, 6, and 16–20 post-infection, and at contemporaneous time points in the uninfected macaques to control for any effect of repeated biopsy on liver immunology. Liver tissue was also acquired at the necropsy time point, which was planned at Week 32 post-infection and occurred earlier out of necessity for some SIV-infected macaques that developed symptoms of simian AIDS (sAIDS).

Additional samples from these macaques were previously used to evaluate liver pathology and metabolic changes resulting from SIV infection (32). The baseline timepoint (Week −4 biopsy) exhibited little to no MX1 protein expression (**Fig 1A-1B and S1A Fig**). In addition, liver biopsies from the uninfected macaques also exhibited very low levels of MX1 at each of the time points, indicating that obtaining liver biopsies alone was not sufficient to elevate MX1 expression (**Fig 1A-1B**). The levels of liver MX1 increased by Week 2 post-infection and remained elevated throughout the study (**Fig 1A-1B**). At Week 2, MX1 staining was predominantly within CD68+ cells (monocyte/macrophage myeloid cells) in the sinusoids (**S1A-1B Fig**), as well as hepatocytes (visualized as cells lacking CD68 staining in the images) (**S1A-1B Fig**). While CD68+ cell-associated MX1 was consistently present throughout infection, hepatocyte MX1 increased in some livers (e.g., RM103), diminished in some (e.g., RM107) and persisted in others (e.g., RM108) over the 32-week time course (**S1A-1D Fig**). Interestingly, the pattern of MX1 expression in hepatocytes became more punctate later in infection, with punctate MX1 in hepatocytes clearly observable at the time of necropsy (**S1C-1D Fig**). These data indicate that MX1 is expressed in the liver predominantly by CD68+ monocyte/macrophage cells and hepatocytes during the first 32 weeks of SIV infection.

**Figure 1.**
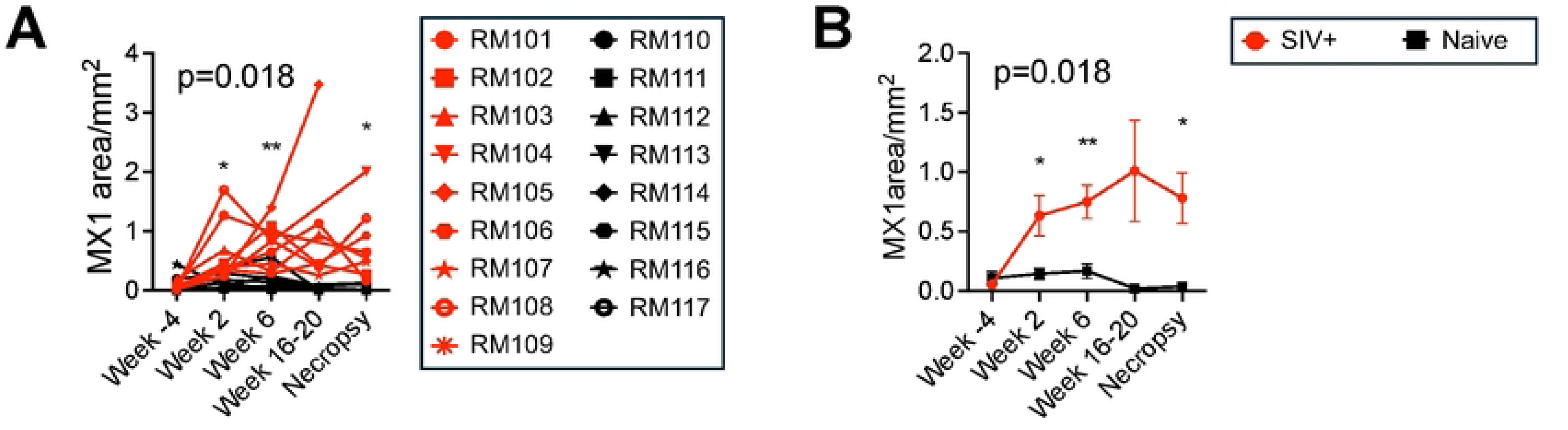
MX1 expression is induced in the SIV infected liver during acute infection and persists over time. FFPE macaque liver biopsies were evaluated by immunofluorescence microscopy to measure expression of the ISG MX1. **(A)** Quantification of MX1 was performed using ImageJ software based on the total area occupied by MX1 signal within a full tissue scan (200x) of the liver section. Each red symbol represents an SIV-infected animal, and each black symbol represents a Naïve-uninfected animal. **(B)** Mean with standard error (SEM) for MX1 signal is shown for the SIV-infected (red) and Naïve-uninfected (black) groups over time.

We previously published that microvesicular steatosis and liver metabolic alterations are hallmarks of acute SIV infection in the liver (32). However, despite the expression of MX1 by hepatocytes throughout infection, no correlation was evident between the percent of hepatocytes with microvesicular (or total including macrovesicular) steatosis and MX1 at Week 2 post-infection (data not shown). Neither was there a clear relationship between the extent of MX1 expressed by hepatocytes at Week 2 and other markers of hepatocyte stress at that time point, such as aminotransferase or cholesterol levels in serum ((32) and data not shown).

However, macaques with more MX1 in hepatocytes at necropsy (e.g., RM103 and RM108, **S1 Fig**) were those that progressed to simian AIDS before Week 32 (32).

Macrophages are crucial in coordinating the hepatic stress response in part through IFN- 1 signaling, and in earlier work, we found increases in both ISGs and CD68+ cells in the livers of chronically SIV-infected macaques compared to uninfected macaques (31). Microscopic analysis determined that the MX1 expression throughout the time course correlated strongly with the number of CD68+ myeloid cells (**Fig 2A**), further implicating these cells in the IFN response. Colocalization analysis exposed an increasing proportion of MX1 signal coming from the CD68+ cells in SIV-infected macaques during acute and early chronic infection followed by a decline in colocalization at the necropsy time point during later chronic infection (percent mean ± standard deviation (SD): 5.29±9.55 Week −4, 7.55±4.31 Week 2, 12.48±6.59 Week 6, 12.98±5.82 Week 16–20, 6.88±3.81 Necropsy), consistent with the pronounced hepatocyte-derived MX1 in many macaques at necropsy (**Fig 2B** and **S1 Fig**). Interestingly, the amount of MX1 signal attributed to CD68+ cells correlated inversely with the quantity of SIV DNA in the liver at contemporaneous time points (**Fig 2C**), suggesting that monocyte/macrophage cells are associated with a reduced SIV burden within the liver.

**Figure 2.**
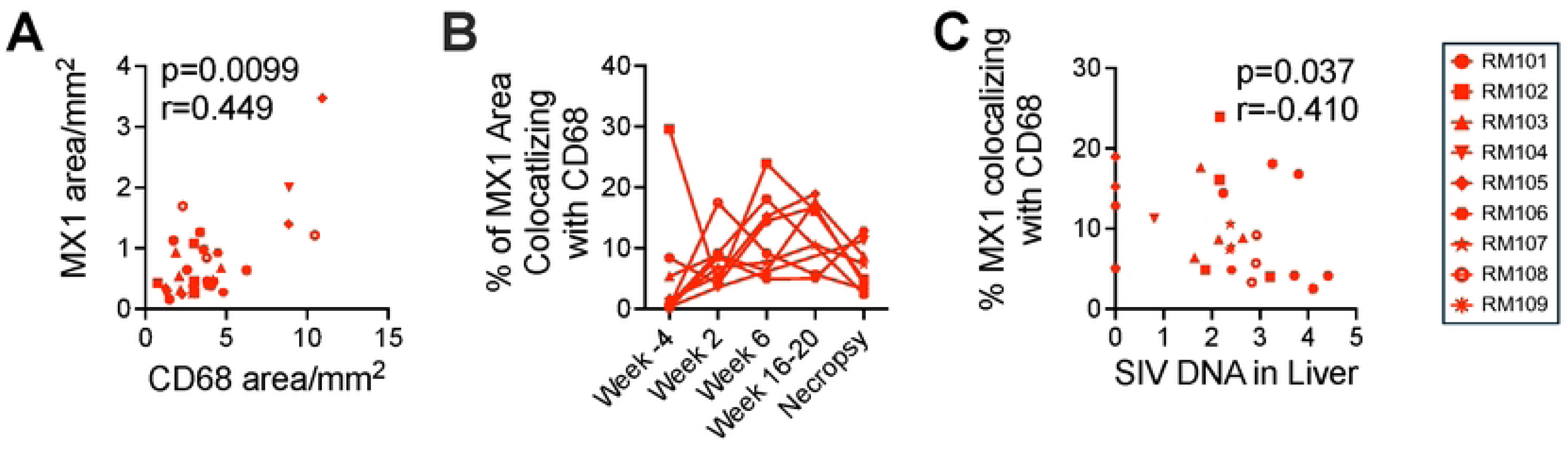
MX1 expression is associated with with CD68+ cells in the SIV-infected liver. **(A)** Immunofluorescence microscopy for MX1 expression was correlated to the expression of CD68 (monocyte/ macrophages) in the liver for each SIV-infected macaque at each biopsy time point. **(B)** The proportion of total MX1 expression derived from monocyte/macrophage cells at each time point by the colocalization of MX1 with CD68 was determined using ImageJ. **(C)** The proportion of MX1 colocalizing with CD68 in each liver biopsy was correlated to the quantity of SIV DNA (copies/million cells) in another liver biopsy at the contemporaneous time point. The quantities of SIV DNA in the livers of these macaques was previously published (32).

### CD206 expression defines monocyte-derived macrophages (MdMs) and Kupffer cells (KCs) in the healthy macaque liver

Myeloid subset identification by flow cytometry was undertaken on freshly isolated liver biopsies based on a previously described flow cytometric gating strategy (33–39). Total liver cells were gated for CD45+ Single Live CD3− CD8− CD20− cells, and within these, myeloid cells were defined as CD11b+ HLA-DR+ (**S2A Fig: 1-6**). The identification of a correlation between the levels of CD11b+ HLA-DR+ cells with those expressing CD68 by flow cytometry (**Fig 3A**) as well as microscopy (**Fig 3B**) provided validation for this approach to identify liver myeloid cells. Furthermore, only a very small percentage of these cells had either a myeloid DC (CD11c+) or plasmacytoid DC (CD123+) phenotype (<0.5% of the cells, and less than 5% of the CD14− cells, **S2B Fig**). Based on these findings, we used CD68+ to define monocytes/macrophages by immunofluorescence microscopy, and CD11b+HLA-DR+ to define them by flow cytometry in the macaque liver.

**Figure 3.**
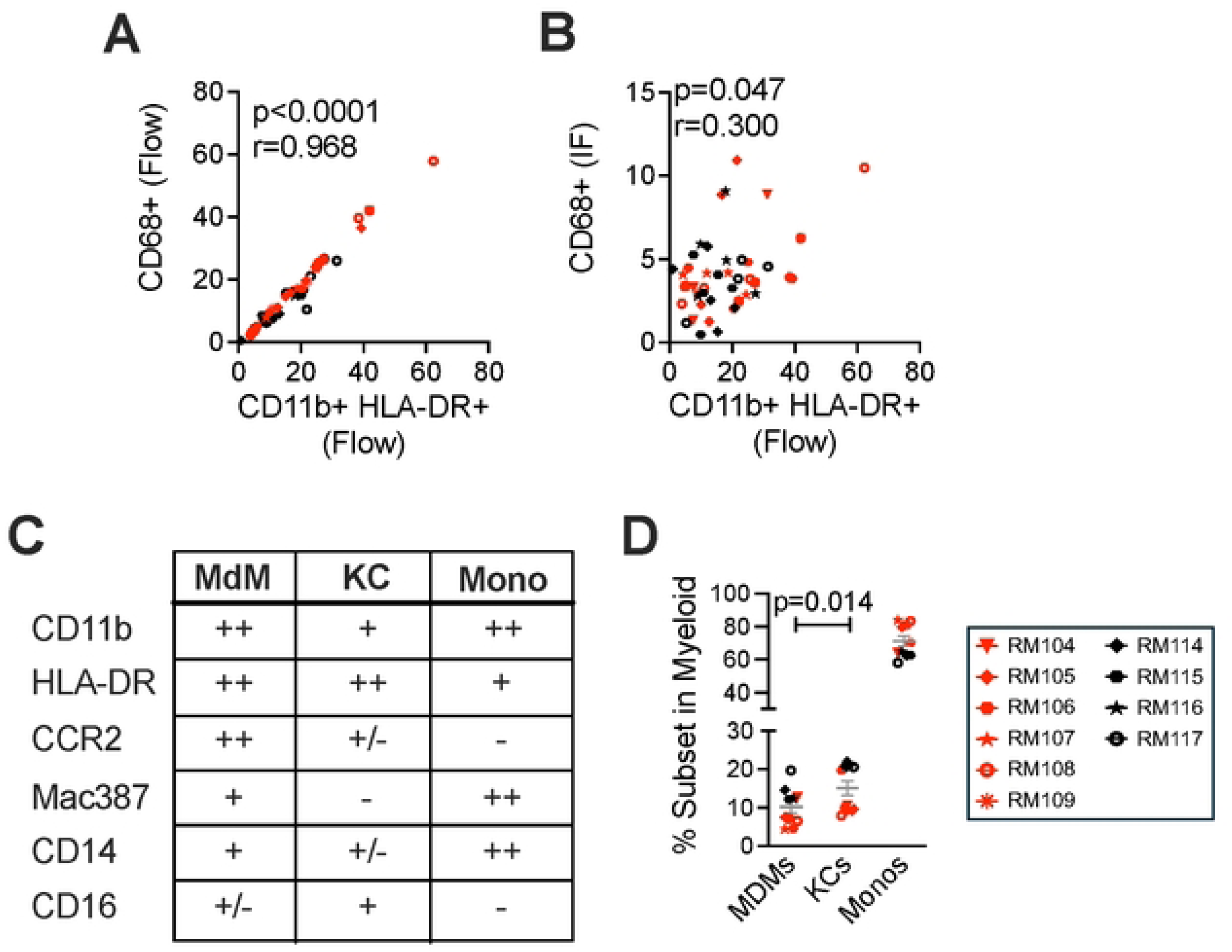
CD163 and CD206 define monocyte/macrophage subsets in the macaque liver. Liver myeloid cell subsets were evaluated in 10 macaques including SIV+ (red) and Naïve- uninfected (black) at each of the study time points. **(A)** The proportion of CD45+ Single Live cells expressing CD68 identified by flow cytometry was correlated with the proportion of CD45+ Single Live CD3- CD8- CD20- cells expressing CD11b and HLA-DR identified by flow cytometry at the same time point. **(B)** The proportion of CD45+ Single Live cells expressing CD68 identified by immunofluorescence microscopy (IF) was correlated with the proportion of CD45+ Single Live CD3- CD8- CD20- cells expressing CD11b and HLA-DR identified by flow cytometry at the same time point. **(C)** Phenotypic markers associated with myeloid cells were evaluated by flow cytometry in uninfected MdM (CD163+CD206−), KC (CD163+CD206+), and Monocyte (Mono) (CD163−CD206−) CD45+ Single Live CD3− CD8− CD20− CD11b+ HLA-DR+ myeloid cells at baseline. **(D)** The proportion of CD11b+ HLA-DR+ myeloid cells defined as MdMs, KCs and Monos by flow cytometry was quantified for all 10 macaques at baseline.

The CD11b+HLA-DR+ cells were then subdivided into populations based on expression of CD163 and CD206 as reported in the literature (33–39): CD163+CD206−, CD163+CD206+, and CD163−CD206− populations (**S2 Fig: 7**). Applying published characterizations of recruited vs resident macrophages in the macaque lung (33) and descriptions of human liver macrophages (40), we identified recruited monocyte-derived macrophages (MdMs, CD163+CD206−), resident Kupffer cells (KCs, CD163+CD206+), and a population comprised predominantly of monocytes (Monos, CD163−CD206−). We defined the cells further by examining expression of monocyte/macrophage markers commonly used across studies in humans, non-human primates, and mice (33, 40–42) as well as markers of recruitment and differentiation (**Fig 3C** and **S3 Fig**). Examining cells at baseline (Week ‒4), we observed that KCs expressed lower levels of CD11b than MdMs or Monos, high amounts of HLA-DR, little to no CCR2 or Mac387 as well as low levels of CD14 and some CD16 (**Fig 3C** and **S3 Fig**). MdMs expressed high levels of CD11b and HLA-DR, were more likely to express CCR2 than KCs or Monos, expressed some Mac387 and CD14 and little CD16 (**Fig 3C** and **S3 Fig**). Monos expressed high levels of CD11b, lower levels of HLA-DR than KCs or MdMs, hardly any CCR2 or CD16 but almost uniformly expressed Mac387 and CD14 (**Fig 3C** and **S3 Fig**). In line with data obtained from rhesus macaque tissues (31, 33, 36, 39), humans (42–44), and mice (45, 46), this expression profile provides clear evidence that the two CD163+ populations (MdMs and KCs) are indeed distinct subsets of liver macrophages. In contrast, CD163− myeloid cells were almost all Mac387+ CCR2− CD14+ and CD16− indicating that they are likely a circulating monocyte population and not a resident or recruited macrophage subset (and not DCs) (**Fig 3C**, **S2B Fig** and **S3 Fig**). Assessment of the proportional representation of these cells at baseline determined that MdMs represented 10.2% (±4.9) (mean ± SD), KCs 15.1% (±6.1), and Monos 71% (±10.0%) of liver myeloid cells (**Fig 3D**). KCs were present at a significantly higher frequency than MdMs (**Fig 3D**).

### KCs accumulate in the SIV+ liver in association with different stimuli and more MX1 than MdMs

We quantified how the frequencies of monocyte/macrophage cells changed over time during SIV infection in livers obtained from 6 SIV+ and 4 naïve-uninfected macaques. While the frequency of liver myeloid cells (CD11b+ HLA-DR+ CD3− CD8− CD20−) was not observed to significantly change during the first 32 weeks of SIV infection, a trend towards an increase (compared to the Week −4 baseline) could be observed (**Fig 4A** and **4C**). Interestingly, the three macaques with the highest proportion of liver CD11b+ HLA-DR+ cells at the necropsy were the ones to develop signs of sAIDS and were euthanized prior to the end of the 32-week study timeline (Week 12 for RM104, Week 27 for RM108, and Week 9 for RM109). Assessment of the distinct myeloid subtypes within the liver determined that MdMs and KCs were both significantly increased at Week 6 post-infection (**Fig 4B** and **4D**), a time that coincides with the resolution of acute plasma viremia and a drop in the quantity of SIV DNA in the liver in these animals (32).

**Figure 4.**
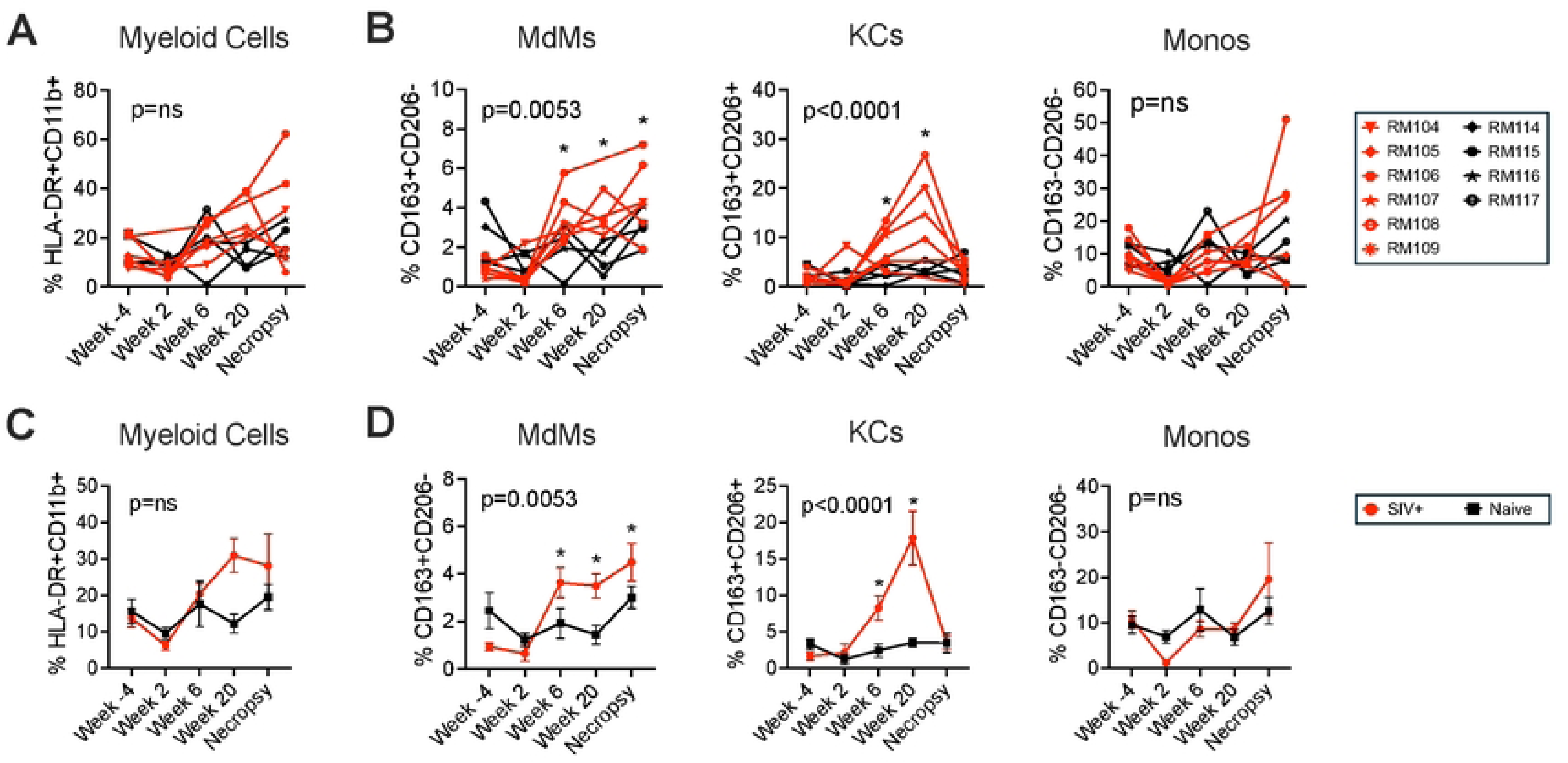
Hepatic KC and MdM frequencies are elevated at multiple time points following SIV infection. **(A)** Longitudinal quantification of the percent of CD45+ Single Live CD3− CD8− CD20− liver cells expressing the myeloid cell-defining markers HLA-DR and CD11b within SIV+ (red) and Naïve-uninfected (black) macaque livers. **(B)** Longitudinal quantification of the percent of myeloid cells (as defined in (A)) defined as MdMs (CD163+CD206−, left), KCs (CD163+CD206+, middle), or Monos (CD163−CD206−, right). Means with standard error are shown for the percent of myeloid cells **(C)**, as well as for MdMs, KCs and Monos **(D)** within the SIV+ as well as the Naïve-uninfected groups.

Elevated levels were also observed at Week 16-20, as well as at necropsy in the MdMs. In contrast, the frequency of Monos did not change significantly throughout the study, although there was a trend toward increased Monos at necropsy among some SIV-infected macaques. Interestingly, the three macaques with the highest frequency of myeloid cells and sAIDS were the same three with the highest proportion of both MdMs and Monos at necropsy while KCs were not elevated in those macaques (**Fig 4B**).

To evaluate factors associated with the changes in myeloid cell subtypes we carried out a correlative assessment of the cell subset frequencies with measured quantities of SIV DNA and 16s bacterial DNA in the livers of the macaques. Evaluating time points in which the changes in myeloid cell frequencies could be observed in the liver (Week 6 through necropsy) we determined that the frequency of MdMs in the liver correlated with the amount of SIV DNA in the liver at the contemporaneous time points (**Fig 5A left**). By contrast, the frequency of KCs over the same time span did not correlate with liver SIV DNA (**Fig 5A right**) (or with SIV RNA in the plasma (32)). Quantities of 16s bacterial DNA were significantly increased in the SIV- infected macaques at Week 16-20 as well as at necropsy (**S4 Fig**). Evaluating the association between 16s bacterial DNA and myeloid subsets at the Weeks 16–20 and necropsy time points in SIV-infected and uninfected macaques identified a significant correlation for both macrophage subsets (**Fig 5B**). Interestingly, while MdMs correlated positively with 16s DNA copies, KCs correlated inversely, suggesting that bacterial products may be influencing MdM frequency, but are likely not driving the accumulation of KCs. While we included uninfected macaques in the analysis, the correlation for KCs held even when uninfected macaques were excluded (SIV- infected macaques only: p=0.015 inverse correlation for KCs). In a previous study, we observed that acute SIV infection was associated with transient hepatocyte damage, which was identified histologically by the presence of the microvesicular form of steatosis, and molecularly by increased expression of the beta-oxidation gene *PPARA* (32). Here the proportion of hepatocytes exhibiting microvesicular steatosis at Week 2 (32) correlated directly with the frequency of KCs, but not MdMs, at the Weeks 6, 16-20, and 32/necropsy time points (p=0.02) (**Fig 5C**). This finding identifies an association between accumulation of KCs and the steatosis that occurs during the acute phase of the infection and suggests that steatosis may be a trigger for KC accumulation. For comparison, we performed similar correlative analyses on liver Monos, with which we found a significant positive correlation with 16s DNA but neither SIV liver DNA nor microvesicular steatosis (**S5 Fig**). These data suggest that, like MdMs, the frequency of Monos infiltrating the liver may be influenced by the presence of bacterial components.

**Figure 5.**
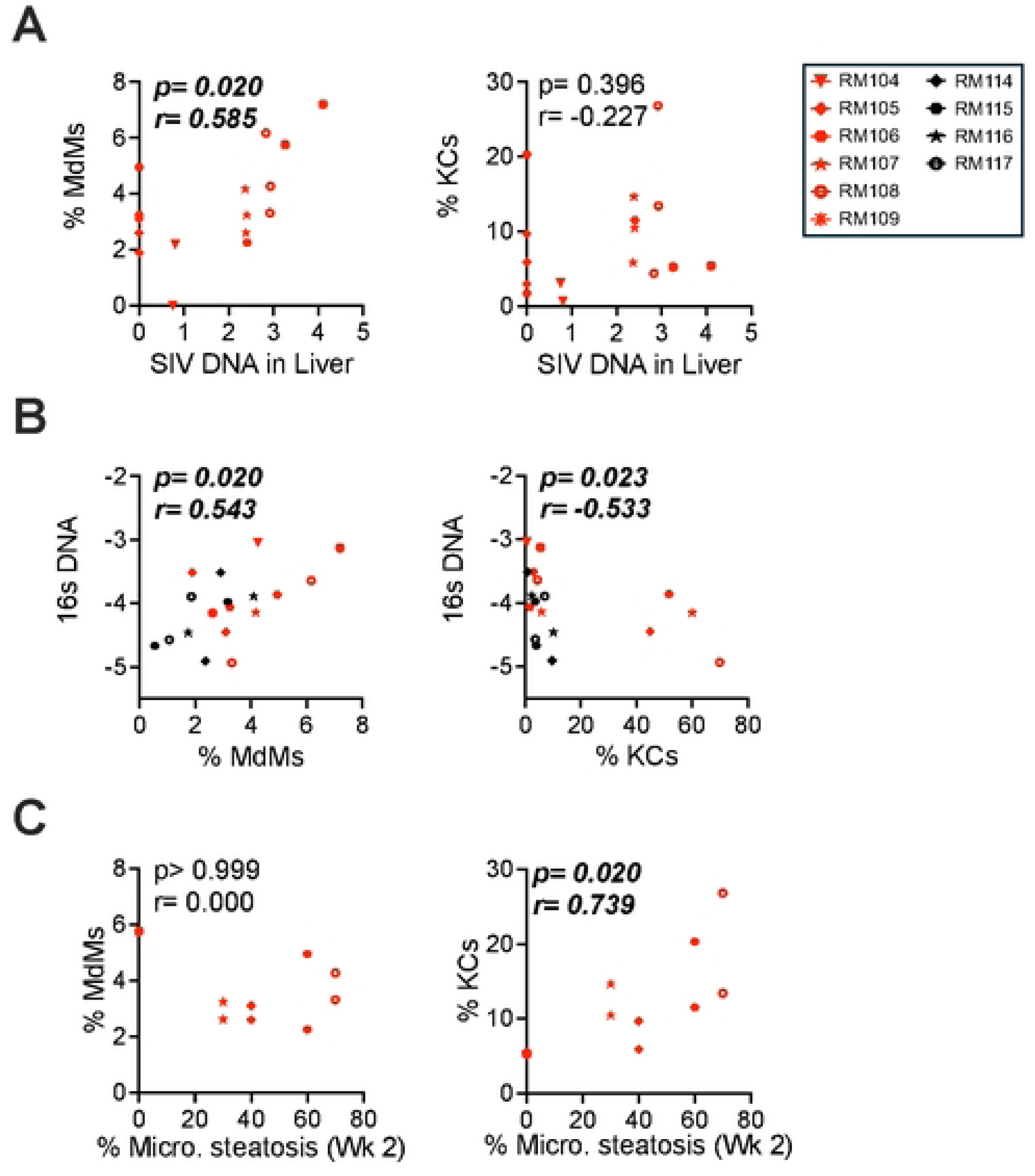
KCs and MdMs diverge in their relationship to SIV load, bacterial infiltration and microvesicular steatosis in the SIV-infected macaque liver. Correlations are shown between the proportion of the indicated cell type in the liver (MdMs left, KCs right) and key parameters of infection and disease outcome. **(A)** Correlation between MdM and KC frequency with SIV liver DNA (copies/million cells) for SIV-infected macaques at Weeks 6, 16-20, and necropsy. **(B)** Correlation between MdM and KC frequency with 16s rRNA DNA copy number (copies/100 ng DNA) in the liver for each SIV-infected and Naïve-uninfected macaque at Week 16-20 and Necropsy. **(C)** Correlation between MdM and KC frequency at Weeks 6 and 20 with the proportion of hepatocytes exhibiting microvesicular steatosis at Week 2. In (A) through (C), significant p-values are italicized and bolded.

We then quantified the expression of MX1 within liver MdMs, KCs and Monos using flow cytometry on cryopreserved cells from necropsy to determine the specific contribution of each subset to expression of this ISG. We determined that the expression could be divided into a MX1^low^ and MX1^high^ phenotype, and the KC population contained significantly more MX1^high^ cells than either the MdM or Mono populations (**Fig 6A**). Furthermore, assessing the geometric mean fluorescence intensity (GMFI) determined that the level of expression per cell within the MX1^high^ populations was significantly higher for KCs than MdMs or Monos (**Fig 6B**). These findings identify the KCs as a major producer of MX1 during chronic SIV infection when compared to the other myeloid subsets.

**Figure 6.**
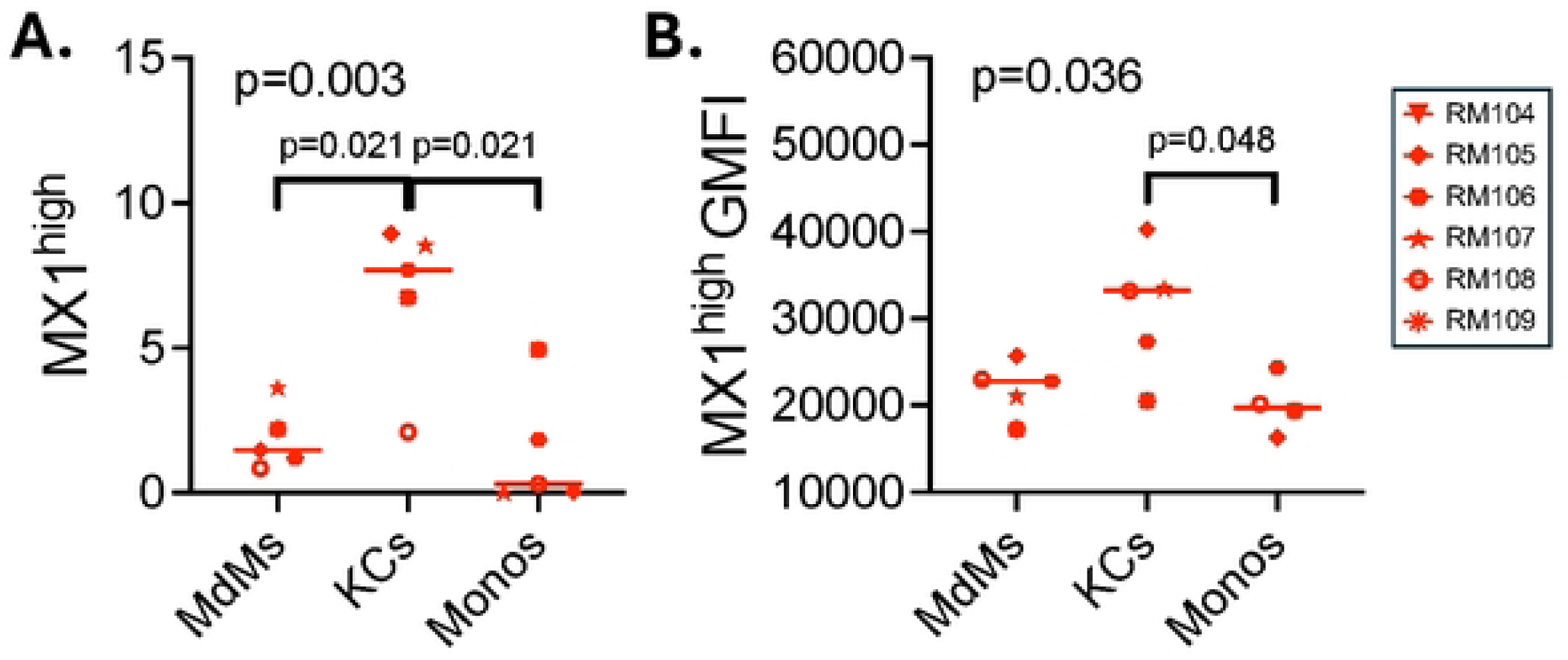
KCs contribute more to MX1 production than MdMs during chronic infection. **(A)** Quantification of flow cytometry to detect MX1 expression within myeloid cells identified the percentage of liver MdMs, KCs, and Monos from SIV infected macaques at necropsy that express high levels of MX1. (**B**) The geometric mean fluorescence intensity (GMFI) of the MX1 signal for the MX1 high population is shown for the liver MdM, KC, and Mono populations.

### KCs and MdMs accumulate in distinct regions of the liver lobule during SIV infection

Liver function is tightly controlled spatially within different zones of the liver lobule (47). Microscopy was utilized to determine the locations of the myeloid subsets within the livers of the SIV+ macaques during acute (Week 2) and chronic (necropsy/Week 32) infection. CD68 was used as a marker of all monocyte/macrophage cells, CD163 for MdMs and KCs and CD206 for KCs and LSECs. Consistent with the flow cytometry data, there were no significant differences between the SIV-infected and uninfected macaque livers at Week 2 in the expression of CD68, CD163, or CD206 (**Fig 7A left**). In contrast, expression of CD68 and CD163 was significantly increased in SIV-infected livers at necropsy (**Fig 7A right**). A representative image detailing the expression of these markers by immunofluorescence microscopy is presented for the liver of an SIV+ macaque obtained at necropsy (**Fig 7B**).

**Figure 7.**
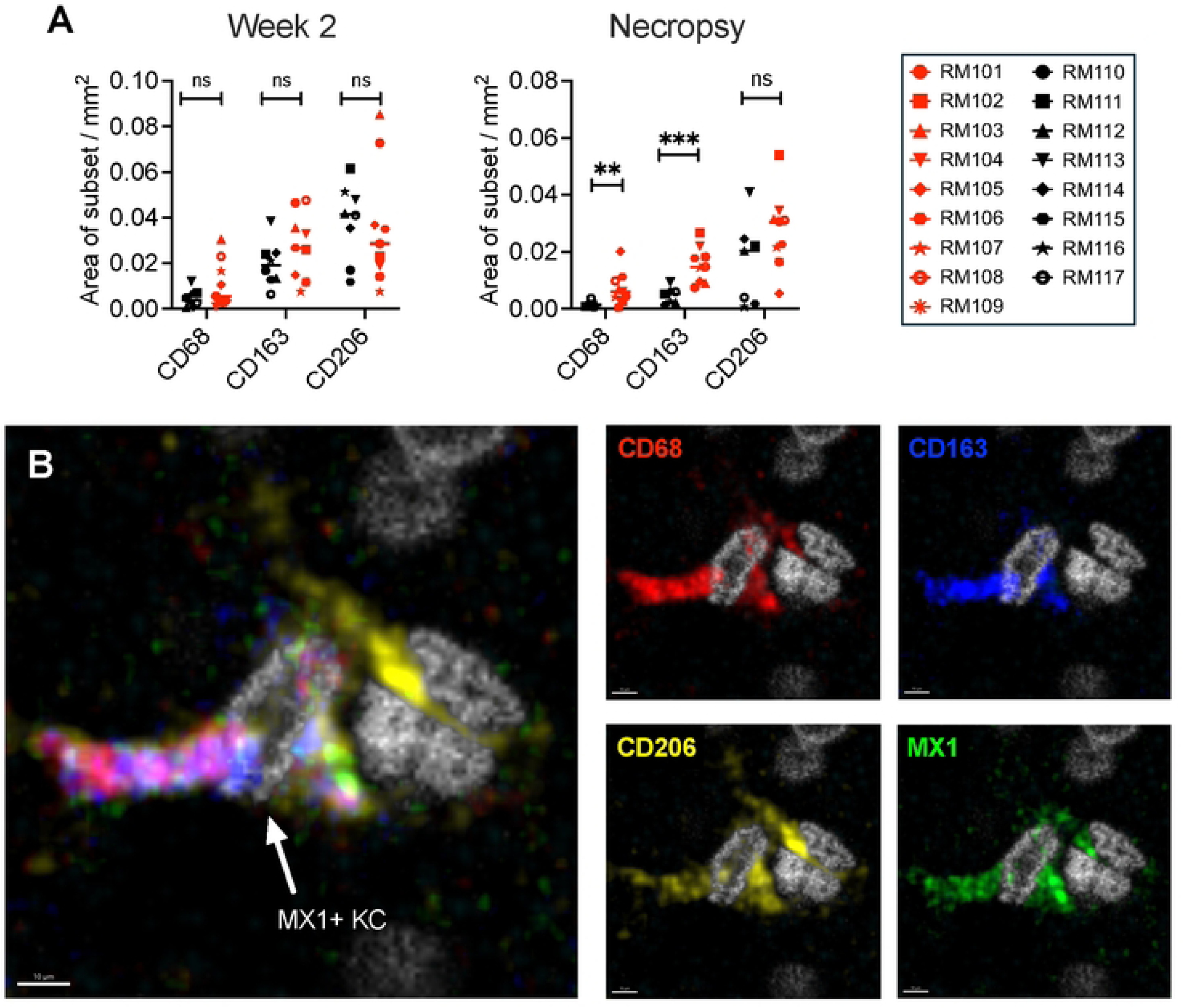
Immunofluorescence microscopy for CD68, CD163 and CD206 identifies monocyte/macrophage populations in their native locations in liver tissue. **(A)** Quantification of the area occupied by monocyte/macrophage associated proteins CD68, CD163 and CD206 within the livers of SIV-infected (red) and uninfected Naïve (black) macaques at week 2 post-infection (left). Quantification of the area occupied by CD68, CD163 and CD206 within the livers of SIV-infected (red) and uninfected Naïve (black) macaques at the Necropsy time point (right). **(B)** A representative confocal image utilized for this quantification is presented for an SIV-infected macaque liver with the expression of CD68 (red), CD163 (blue), CD206 (yellow), and MX1 (green) along with nuclei (NucSpot DNA stain, gray) depicted. The 4- color overlay is shown at the left and single-color images to the right. An MX1+ KC is indicated by the arrow (CD68+CD163+CD206+MX1+).

To clarify macrophage locations during early (Week 2) vs late (necropsy/Week 32) infection and identify possible zonal differences in production of MX1, a high parameter antibody staining panel was used that detects expression of CD68, CD163, CD206 and MX1 on the same cells within different zones of the liver lobule by microscopy (**Fig 8A** and **8B**). We marked the periportal zone by hepatocytes expressing arginosuccinate synthetase 1 (ASS1) and the centrilobular zone by hepatocytes expressing glutamine synthetase (GS) as we did previously to observe T cell location (32). This strategy clearly identified MdMs (CD68+CD163+CD206−) and KCs (CD68+CD163+CD206+) (**Fig 8B**). It also revealed the presence of CD163−CD206+ cells, which are likely LSECs based on their location, morphology and bright CD206 expression (48) (**Fig 8A**). MdMs accumulated in the periportal zone of the lobule with the increase evident at necropsy (p=0.04 vs. centrilobular zone) but not Week 2 (**Fig 8C left**). In contrast, KCs accumulated within the centrilobular zone and declined in the periportal zone with the shift already beginning by Week 2 and significant at necropsy (p=0.002) (**Fig 8C middle**). Finally, an increase in CD163−CD206+ putative LSECs was detected within both zones, especially near the central veins, at necropsy (p=0.048 necropsy vs. week 2) (**Fig 8C right**). While KCs and MdMs both expressed MX1 at necropsy, KCs expressed MX1 earlier than MdMs, especially in the centrilobular zone (p=0.036 KC vs MdM week 2) and persisted in having high MX1 expression at necropsy (**Fig 8D left, middle**). Interestingly, the putative LSECs also expressed MX1, with expression peaking at Week 2 and then declining by necropsy but remaining elevated compared to uninfected macaques (**Fig 8D right**). These findings provide insights regarding the location of the MdMs, KCs and LSECs and provide additional microscopic evidence that SIV infection is associated with accumulation of these cell subsets in the liver with differing contribution to MX1 production.

**Figure 8.**
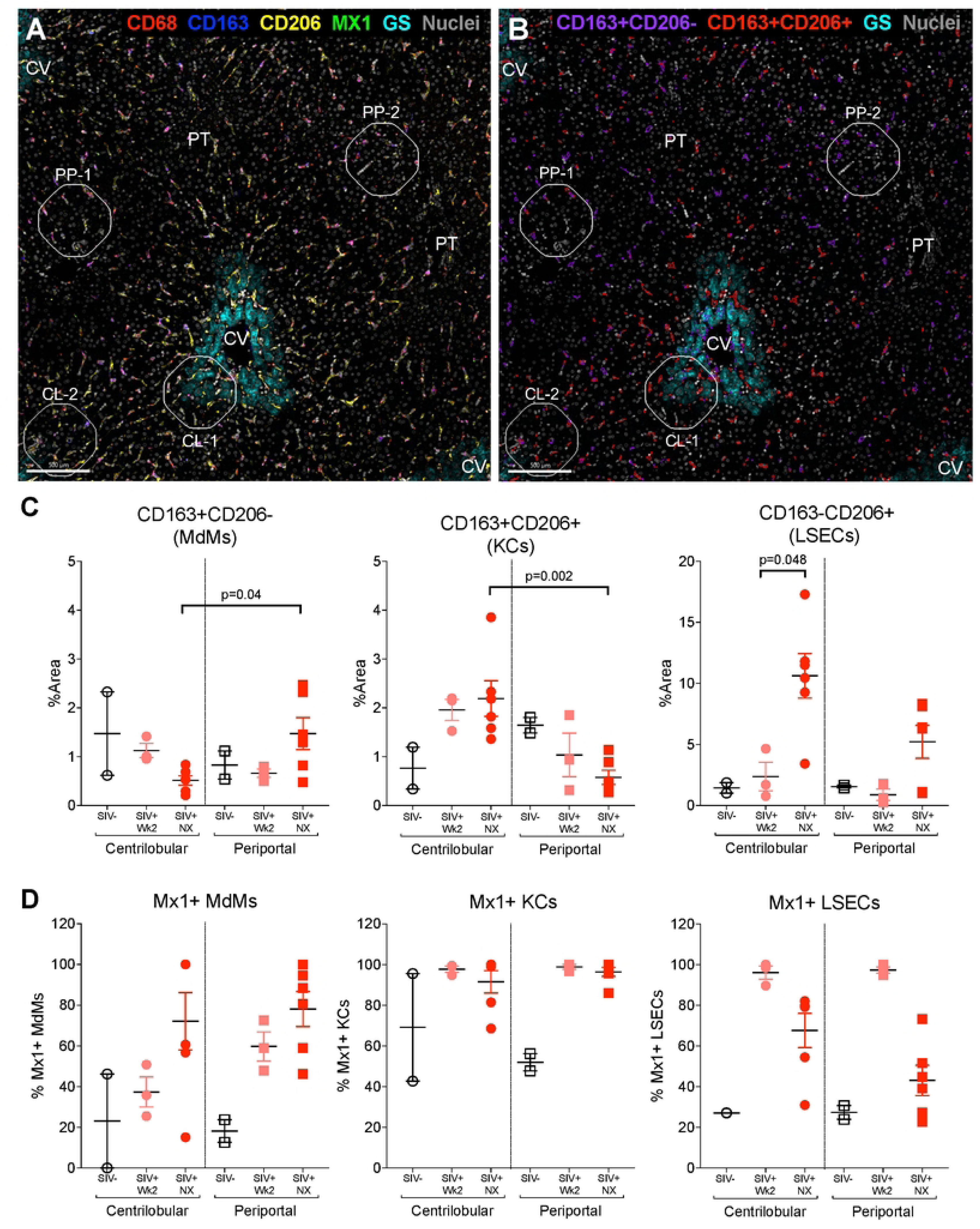
KCs shift towards the centrilobular zone during SIV infection. **(A)** Confocal microscopy image at 200X of an SIV-infected macaque liver at necropsy with central veins (CV) and portal tracts (PT) noted. The CV is demarcated by expression of GS (cyan) in pericentral hepatocytes. Expression of CD68 (red), CD163 (blue), CD206 (yellow), MX1 (green), and nuclei (gray) are also shown. **(B)** Imaris surfaces demarcating CD163+CD206− MdMs (purple) and CD163+CD206+ KCs (red) along with GS and nuclear staining in a tissue from an SIV-infected macaque at necropsy. In **(A, B)** ROIs used for analysis are noted. **(C)** Quantification of CD163+CD206− MdM areas (left), CD163+CD206+ KC areas (middle), and CD163-CD206+ LSEC areas (right) from 2-4 ROIs from 1-3 macaques in each group. The y-axis indicates the percent of the ROI area occupied by the indicated fluorescent signals. **(D)** Quantification of MdM (left), KC (middle), and LSEC (right) areas co-expressing MX1. The y-axis indicates the percent of the MdM, KC, or LSEC fluorescent area colocalizing within 0.1μm with MX1 staining.

## Discussion

Liver disease is a major contributor to morbidity and mortality in PWH, even with suppressive cART (5, 6, 8, 10, 49). Here we used liver biopsies obtained from SIV-infected rhesus macaques to evaluate changes in the hepatic IFN-1 response during the acute and chronic phases of infection. Cell phenotyping, spatial mapping and correlation analyses were applied to examine the temporal expression of a canonical ISG (MX1), the contributions of discrete monocyte/macrophage subsets to MX1 expression, the spatial distribution of macrophage subsets within the liver lobule, and factors associated with temporal and spatial changes in macrophage populations. Our findings provide insights into the factors that may promote elicitation of the IFN-1 response within the liver following SIV infection and the cell subsets that contribute to this response.

Previously, we determined that hepatocytes are affected by SIV already at Week 2 post-infection: they exhibit microvesicular steatosis, the liver expresses elevated levels of *PPARA* mRNA, and the serum carries higher concentrations of aminotransferases and cholesterol (32). Here we more deeply characterized the Week 2 biopsy time point to inform on the immune environment associated with the hepatocyte effects during acute SIV infection. Microscopy revealed increased MX1 expression in the liver at Week 2, including within hepatocytes (**Fig 1**). As hepatocytes are the predominant cell type in the liver, even a small increase in the expression of ISGs per cell (as shown in **S1 Fig**) has the potential to have a large effect on overall expression of the protein in the tissue. Induction of ISGs in hepatocytes is well-documented during infection with Hepatitis B and C viruses, which replicate within hepatocytes directly (50) and could potentially result from the binding of CCR5 on hepatocytes to gp120, which has been observed *in vitro* even though hepatocytes are not productively infected by HIV or SIV *in vivo* (51).

Importantly, the intensity of MX1 staining in hepatocytes at Week 2 was less than that observed in CD68+ myeloid cells (**S1 Fig**). While the frequencies of the three major myeloid subsets (MdMs, KCs and Monos) were not increased compared to baseline at this time point (**Fig 4** and **Fig 7**), MX1 expression was elevated within the resident KCs and also within CD163–CD206+ (LSEC) cells, implicating these two liver-resident cell types as early producers of MX1 along with hepatocytes prior to monocyte/macrophage infiltration (**Fig 8**). We did not expect to identify a role for LSECs in the response to SIV in the liver; however, it is not surprising as these endothelial cells that line the hepatic sinusoids and express high levels of scavenger receptors (including CD206) are involved in endocytosis of blood-borne pathogens and PAMPs (52–54). To inform on the spatial dynamics of the response, we evaluated monocyte/macrophage and LSEC positions within the liver lobule. Macrophage skewing towards the periportal zone is a recognized characteristic of the healthy liver while altered macrophage zonation is associated with bacterial translocation (55). Although we observed a trending shift in KC location towards the centrilobular zone at Week 2 (**Fig 8**), we did not detect heightened levels of bacterial 16s DNA in the liver at that time point (**S4 Fig**), suggesting that the early difference in zonation is not mediated by bacteria translocating from the gut. Overall, the analysis of Week 2 liver biopsies identified an increase in MX1 expression that was likely produced within KCs, LSECs and hepatocytes. These findings indicate that resident cells in the liver respond to SIV infection through the production of ISGs as early as 2 weeks post SIV- infection.

The chronic phase of SIV infection was evaluated using the liver biopsies obtained at Weeks 6, 16-20 and 32/necropsy. At these time points, we observed increased MX1 expression in four different cell types compared to baseline and uninfected macaque livers: hepatocytes, LSECs, MdMs and KCs (**Fig 6-8** and **S1 Fig**), and we also found increased frequencies of the MdMs and KCs in the livers (**Fig 4**). Although KCs declined from their peak accumulation by necropsy/Week 32 (**Fig 4**), these cells expressed more MX1 than MdMs or Monos at that time point (**Fig 6**) with equivalently high expression by KCs in periportal and centrilobular zones (**Fig 8**). MdMs in both zones also expressed MX1 albeit at lower levels than KCs (**Fig 6** and **Fig 8**). Interestingly, CD206+CD163– LSECs declined in their MX1 expression from Week 2 to necropsy/Week 32 indicating a lower contribution to ISG production later in the infection despite higher levels of bacterial DNA (**Fig 8**). These findings further support that hepatocytes, LSECs, MdMs and KCs each likely play a key role in production of ISGs during SIV infection. However, the KCs were the highest ISG expressors, which identifies them as a potentially critical cell population in the inflammation and immune activation observed in PWH. Similar findings have been reported in the literature on people with cirrhosis and Hepatitis Virus infection wherein human KC-like macrophages (CD206+ HLA-DR^high^) in the liver have been shown to take on a pro-inflammatory M1 phenotype, expressing elevated levels of CD14 and producing TNF-α and GM-CSF (41, 42).

The expression of MX1 and other ISGs is typically the result of PAMP-initiated production of IFN-1 and cell responses through the IFN receptor (IFNAR). Based on our previous studies (30–32), there are two PAMP-containing entities that have the greatest potential to drive IFN-1 production in the SIV-infected macaque liver: 1. SIV, either in the form of free or cell-associated virus, and 2. Bacteria (or bacterial products) translocated into the liver through an SIV-compromised gut barrier. Using correlation analyses to evaluate the relationships of these potential factors to key myeloid cell populations (**Fig 5** and **S5 Fig**), we identified SIV DNA load in the liver as a correlate with MdM frequency at chronic time points. This suggests that SIV may be inducing liver recruitment of MdMs or local expansion of macrophages with an MdM phenotype whereas this was not the case for KCs or monocytes (with which SIV DNA load did not correlate). Evaluating the association between 16S bacterial DNA and myeloid subsets at these post-infection time points identified a significant increase for all three subsets, underscoring a critical role of bacterial components in the hepatic IFN-1 response. Unlike MdMs and Monos, KCs correlated inversely, suggesting that bacterial products are likely not driving the accumulation of these cells. In addition to PAMPs, host-derived damage-associated molecular patterns (DAMPs) including mitochondrial DNA are recognized as triggers of inflammatory responses (56) and we previously detected microvesicular steatosis during acute SIV infection (32). As this form of hepatocyte lipid accumulation is often associated with mitochondrial dysfunction (57), we considered the possibility that early steatosis might trigger KC accumulation and IFN-1 signaling. Indeed, microvesicular steatosis at Week 2 correlated with KC accumulation at the ensuing time points (**Fig 5**), suggesting there may indeed be a role for host DAMPs in the SIV-infected liver in driving IFN-1 signaling and supporting the possibility that the microvesicular steatosis triggers KC MX1 production.

In adults with steatohepatitis, liver fibrosis generally begins around central veins (58), yet previous work in the murine and human liver have demonstrated a polarized localization of macrophages towards the periportal zones and portal tracts during homeostasis (55). We found that not only was the frequency of KCs inversely proportional to the quantity of 16s DNA during chronic infection, but as early as Week 2, the KCs were positionally shifted in the lobules with greater concentration towards the central veins. Published data show that bacterial components are detectable throughout the lobules in the chronically SIV-infected liver (21), potentially representing a cue for macrophage trafficking and response or a result of macrophage movement. As KCs are already shifting spatially from a predominantly periportal localization to a centrilobular localization at Week 2, it is likely that microbial translocation is not the driver of the KC migration, but that KC migration may allow microbes to migrate deeper into the lobules, gaining easier access to the systemic circulation. Indeed, in mice lacking periportal KC localization, microbes were better able to spread through the liver lobule and access the peripheral blood (55).

Taken together, these findings provide evidence for a role of multiple cell types in the IFN-1 response within the liver following SIV infection. KCs, which correlate with early hepatocyte steatosis, exhibited the greatest capacity for ISG expression and demonstrate a potent IFN-1 response in the liver even at 2 weeks post-infection. We hypothesize that with centrilobular zone accumulation of KCs, the IFN-1 response by KCs may set the stage for the initiation of central vein fibrosis and hepatocyte damage in later stages of the infection. Specific modulation of KC function, myeloid cell accumulation and IFN-1 response may prove valuable strategies to limit liver disease in PWH.

## Materials and Methods

### Ethics statement

The animal study from which the liver samples used herein were derived was approved by the Institutional Animal Care and Use Committee (IACUC protocol # IP00001485) of the Oregon National Primate Research Center (ONPRC, Beaverton, OR, USA). All rhesus macaques involved in the study were cared for according to the laws, regulations, and guidelines set forth by the United States Department of Agriculture, Institute for Laboratory Animal Research, Public Health Service, National Research Council, Centers for Disease Control, the Weatherall Report titled “The use of non-human primates in research”, and the Association for Assessment and Accreditation of Laboratory Animal Care (AAALAC) International. Nutritional plans used by the ONPRC consisted of standard monkey chow supplemented with a variety of fruits and vegetables as part of the environmental enrichment program established by the Behavioral Management Unit. Enrichment was distributed and overseen by veterinary staff with animals having access to more than one category of enrichment. SIV-infected macaques were housed in individual but adjoining cages allowing for social interactions. Primate health was observed daily by trained staff. All efforts were made to minimize suffering using minimally invasive procedures, anesthetics and analgesics when deemed appropriate by veterinary staff. Animals were painlessly euthanized via sedation with ketamine hydrochloride injection followed by an intravenous barbiturate overdose following the recommendations of the panel of euthanasia of the American Veterinary Medical Association.

### Study design and liver biopsy processing

A total of 17 adult Indian rhesus macaques were enrolled in a study to investigate the influence of SIV infection on liver health over time (32). In that study, nine macaques were infected intravenously with SIVmac251 at Week 0, and the other eight macaques served as SIV-naïve uninfected controls. Laparoscopic liver pinch biopsies were obtained as described (32) from each animal in the SIV group four weeks before SIV inoculation (Week −4), as well as at four time points following infection: Weeks 2, 6, 16–20, and Necropsy (approximately Week 32).

Liver biopsies were collected from the SIV-naïve uninfected macaques at the same times. The study was split in two cohorts. SIV-infected macaques RM101-103 and uninfected macaques RM110-113 comprised the first cohort with samples at Week 16 while SIV-infected macaques RM104-109 and uninfected macaques RM114-117 comprised the second cohort with samples at Week 20. At each time point from each macaque, one biopsy was flash-frozen and one was formalin-fixed paraffin-embedded (FFPE) as described (32). In the second cohort, a third biopsy was collected for the isolation of single cells. The biopsy was rinsed in phosphate buffered saline (PBS), minced, and incubated at 37C for 30 minutes with broad spectrum collagenase (NB8 derived from *Clostridium histolydicum*, Nordmark, Uetersen, Germany) prepared in a buffer of Hanks balanced salt solution with calcium and magnesium (HBSS +/+) containing added 1.5% fetal bovine serum (FBS), 2mM calcium chloride, and 5mM HEPES. Following incubation, the cells were passed through a 70μm cell strainer, washed twice with HBSS without calcium/magnesium (HBSS -/-), treated for red blood cell lysis, and washed again before counting in Roswell Park Memorial Institute medium 1640 (RPMI) with 10% FBS. Total recovered liver cells were used immediately for flow cytometry assays or cryopreserved in the vapor phase of liquid nitrogen. At the necropsy, larger pieces of liver tissue were obtained and processed according to the same protocol.

### Immunofluorescence microscopy

Antibodies to CD68, CD163, CD206, MX1, arginosuccinate synthetase 1 (ASS1), and glutamine synthetase (GS) were used to observe hepatocytes and immune cells in their native locations within FFPE liver biopsies by immunofluorescence microscopy. The antibodies are described in detail in **S1 Table**. Some antibodies were used unconjugated with secondary conjugated antibodies; some were purchased unconjugated and conjugated to dyes in house using Biotium Mix n’ Stain kits (San Francisco, CA, USA); and others were purchased conjugated to dyes for direct immunofluorescence. Biopsies were sectioned 5μm thick onto glass slides and the sections were deparaffinized and rehydrated and antigens unmasked as described (32). After washing in TBS-T (0.025% TritonX-100 in Tris buffered saline) and blocking (0.1% bovine serum albumin (BSA) at 1% goat serum in TBS-T), tissues were incubated with primary antibodies overnight at 4C and washed. For staining performed using unconjugated primary antibodies, secondary antibodies were then added for 60 minutes at room temperature in the dark.

Secondary antibodies were goat anti-mouse-AlexaFluor (AF) 488 (1:500), goat anti-rabbit- AF594 (1:500), and goat anti-rabbit AF647 (1:500). Washed slides were mounted in Vectashield hard-set mounting medium with DAPI (Vector Labs, Newark, CA, USA) or were incubated with 10x NucSpot 750 (Biotium) for 15 minutes at room temperature before mounting in Vectashield hard set without DAPI. Slides were imaged on either a Keyence fluorescence microscope using 20x, 40x, or 100x objectives or on a Stellaris confocal microscope using 20x or 60x objectives. Images were analyzed using Fiji and Imaris software. Fiji was used to measure the area occupied by a cell marker (e.g. CD68, MX1) and to quantify the overlap between CD68 and MX1 pixels for images stained with antibodies to CD68 and MX1 and imaged on the Keyence microscope. Imaris was used to analyze high parameter confocal microscopy data acquired on the Stellaris microscope. In Imaris, surfaces were created for CD163, CD206, and MX1 with standard threshold and voxel values. These surfaces were used to create masks for identifying CD163+CD206+, CD163+CD206–, and CD163–CD206+ cells and creating surfaces based on marker expression. MX1 expression associated with each population was determined with upper threshold bound set to 0.1 to identify MX1-expressing objects ≤ 0.1μm from the surface. To identify zone location of the surfaces, regions of interest (ROIs) were then placed on images based on ASS1 and GS expression without macrophage staining visible to reduce bias in ROI selection. ROIs were circular with 20 vertices and sized to collect approximately 100 cells. Cell populations were then quantified in each ROI and reported.

### Flow cytometry

Single cell suspensions of macaque liver (10^6^ cells per macaque per time point) were incubated with viability dye and antibodies to cell surface markers (**S2 Table**) at room temperature for 20 minutes. Cells were washed in PBS/2% FBS, pelleted, and incubated in 1x Fix/Perm solution (eBiosciences, San Diego, CA, USA) for 20 minutes. Cells were then washed in 1x Permeabilization buffer (eBiosciences), incubated with antibodies to intracellular markers at room temperature for 1 hour, and washed again in 1x Perm buffer at PBS/2% FBS. All staining and washing steps were performed protected from light. Immediately after staining, the cells were analyzed on a BD LSRII analyzer. Flow cytometry was always performed using freshly isolated liver cells except for the MX1 staining, which used cryopreserved liver cells from necropsy. All data were analyzed using FlowJo software (TreeStar, version 10). Gates for cell populations were determined using fluorescence minus one (FMO) stained controls.

### 16s rRNA DNA quantitative PCR

DNA extracted from flash-frozen liver biopsies prepared as described (32) was added to qPCR reactions according to the manufacturer’s instructions for the Femto Bacterial DNA Quantification Kit (Zymo Research, Irvine, CA, USA) as previously described (31).

### Statistics

All data obtained in the study were analyzed using a GraphPad Prism version 9 software (San Diego, CA, USA). Longitudinal data were analyzed using a mixed-effects model with Dunnett’s multiple comparison test. Data from three monocyte/macrophage subsets at a single time point in the same animals were compared using a Friedman paired non-parametric ANOVA with Dunn’s multiple comparison correction test. Data from two macrophage subsets at a single time point in the same animals were compared using a Wilcoxon paired non-parametric t-test. Data from two groups of animals were compared using a Mann-Whitney unpaired non-parametric t-test. Correlations were established by identifying the Spearman correlation coefficient across two sets of data. All results were deemed significant for p<0.05.

## ACKNOWLEDGEMENTS

The authors would like to acknowledge the veterinary and support staff at the Oregon National Primate Research Center, and Xuemei Deng in the Seattle Children’s Hospital Anatomic Pathology Department for technical expertise in cutting tissue blocks.

## AUTHOR CONTRIBUTIONS

ND: study design, overseeing, experiments, analysis, writing, editing BIJ: experiments

SB: overseeing, experiments

KAF: experiments

CLS: experiments

SY: experiments

KAM: experiments

YMA: experiments

SSL: experiments

CF: experiments

MF: animals

JVS: animals

BJB: study design, overseeing, funding

DLS: study design, overseeing, funding, editing

## SUPPLEMENTARY TABLES

## SUPPLEMENTARY FIGURES

**Supplementary Figure 1. Assessment of MX1 and CD68 expression within the livers of SIV infected macaques throughout disease course. (A)** The expression of MX1 and CD68 alongside nuclear staining using DAPI is presented within the livers of three representative SIV- infected macaques. Time points include Weeks –4 (baseline), 2, 4, 16-20 and necropsy. **(B)** An example image from Week 2 post-infection reveals MX1 expression predominantly within CD68+ cells (monocytes/macrophages), with some signal present in CD68– cells (presumably hepatocytes). **(C,D)** Two examples of MX1 and CD68 expression at necropsy are shown from two different SIV-infected macaques exhibiting varying degrees of CD68+ and CD68– MX1 staining.

**Supplementary Figure 2. Gating strategy to identify monocyte/macrophage cell subsets in liver single cell suspensions by flow cytometry.** Single cell suspensions of macaque liver were stained with a cocktail of antibodies to identify monocyte/macrophage cell subsets using a gating strategy based on that by Cai, et al (33). **(A)** A representative SIV-infected macaque at week 20 is shown. CD45+ cells were first selected (1) and from those, debris was excluded (2) and single cells were selected (3). Dead cells were excluded (4), and then cells expressing lineage markers CD3, CD8, and CD20 were excluded (5). Cells were selected that expressed CD11b and HLA-DR with both CD11b-intermediate (CD11b^int^) and CD11b-high (CD11b^high^) expressing cells captured (6). CD11b+ HLA-DR+ cells were termed “Myeloid cells”, and these were quantified as a proportion of the CD45+, Non-debris, Single, Live cells, termed “CD45+”. Within the myeloid gate, CD163+CD206–, CD163+CD206+, and CD163– cells were selected and termed monocyte-derived macrophages (MdMs), Kupffer cells (KCs), and monocytes (Monos), respectively (7). These populations were quantified as a proportion of the CD45+ population defined above. **(B)** Within the CD11b+ HLA-DR+ gate, dendritic cells (DCs) were identified by gating CD14+ vs. CD14– cells and divided into CD11c+ myeloid DCs (mDCs) and CD123+ plasmacytoid DCs (pDCs). As expected, the mDC and pDC populations were only present within the CD14– subset, and these cells represented only a small fraction of the myeloid cells in the liver.

**Supplementary Figure 3. Phenotypic analysis to define MdMs, KCs, and Monos at baseline.** An example of the flow cytometry phenotyping strategy used to define MdM, KC, and Mono signatures is depicted for one macaque at baseline in **(A-C)** and quantified for all macaques at baseline in **(D-K)**. **(A)** Histograms displaying the expression intensity (geometric mean fluorescence intensity (GMFI)) of CD11b (left) and HLA-DR (right) within the CD163+CD206– MdM (yellow), CD163+CD206+ KC (pink), and CD163–CD206– Mono (aqua) populations. **(B)** Graphs displaying CCR2-expressing and Mac387-expressing populations within the MdM (left), KC (middle), and Mono (right) subsets of Myeloid cells. **(C)** Graphs displaying CD16-expressing and CD14-expressing populations within the MdM (left), KC (middle), and Mono (right) populations alongside a key (far right) that distinguishes the myeloid cell subsets based on CD14 and CD16 levels. **(D-K)** indicate the quantification of the MdM, KC, and Mono populations across all 10 macaques included in this analysis for **(D)** CD11b GMFI, **(E)** HLA-DR GMFI, **(F)** % Mac387+, **(G)** % CCR2+, **(H)** % CD14+CD16–, **(I)** % CD14+CD16^int^, **(J)** CD14–CD16^int^, **(K)** CD16^high^.

**Supplementary Figure 4. 16s DNA in the liver during 32 weeks of SIV infection.** 16s rRNA DNA was measured in total DNA extracted from liver biopsies using the Femto bacterial DNA quantification kit. The number of copies of 16s DNA in liver biopsies from SIV-infected (red) vs. Naïve (black) macaques is shown to differ across the time course **(A)** and is significant specifically at the Week 16-20 and necropsy time points **(B)**.

**Supplementary Figure 5. Correlation between the frequency of monocytes in the liver and the detected SIV DNA, 16s DNA and microvesicular steatosis.** Correlations are shown between the proportion of Monos in the liver and key parameters of infection and disease outcome as in Figure 5. Significant p-values are italicized and bolded.

**Supplementary Table 1.** Antibodies for immunofluorescence microscopy.

| Target | Clone | Host | Dilution | Source | Color | Strategy |
| --- | --- | --- | --- | --- | --- | --- |
| ASS1 | EPR12398 | rabbit | 1:500 | Abcam | AF488 | Conj. |
| CD68 | EPR20545 | Rabbit | 1:500 | Abcam | AF594 | 1°, 2°, Conj. |
| CD163 | EDHu-1 | Mouse | 1:50 | BioRad | CF405 | In-house conj. |
| CD206 | Polyclonal | Rabbit | 1:750 | Abcam | AF647 | 1°, 2° |
| GS | GT1055 | Mouse | 1:500 | Thermofisher | CF700 | In-house conj. |
| MX1 | CL143 | Mouse | 1:50 | Millipore | AF488 | 1°, 2° |
|  |  |  |  |  | AF594 | 1°, 2° |
|  |  |  |  |  | CF555 | In-house conj. |

**Supplementary Table 2.**
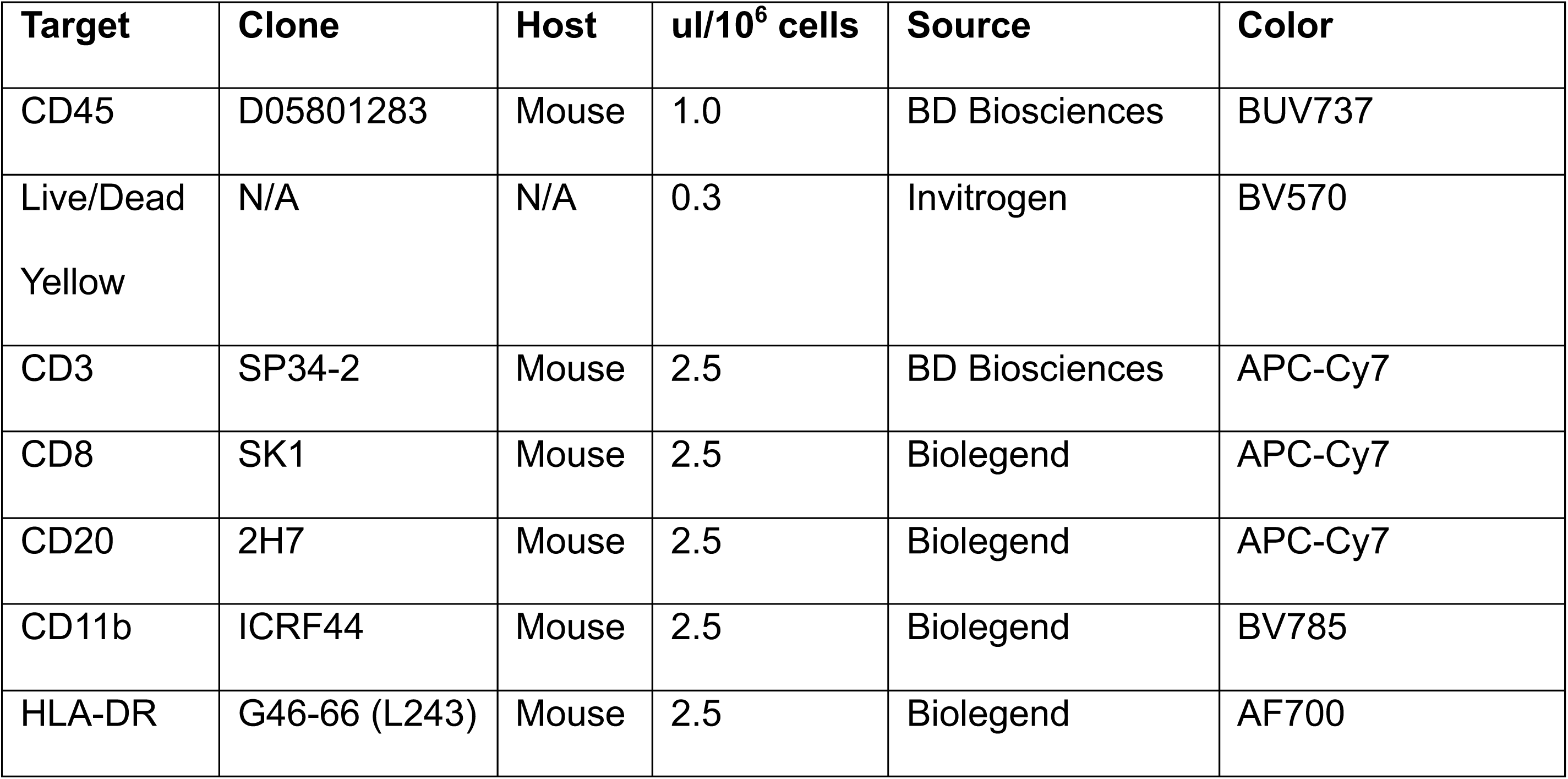
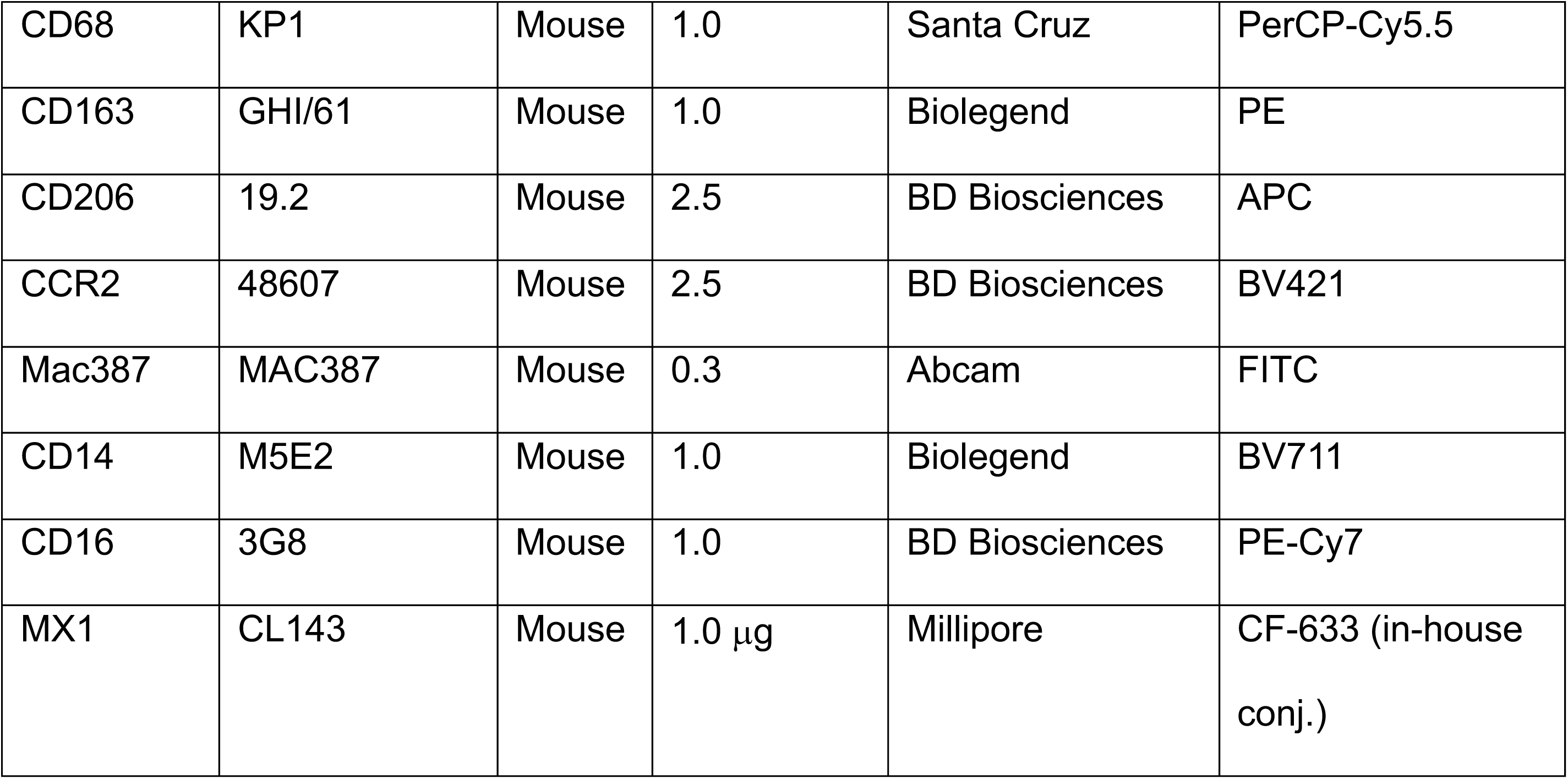
Antibodies and dyes for flow cytometry.

## Notes

### Competing Interest Statement

The authors have declared no competing interest.

